# RNAGEN: A generative adversarial network-based model to generate synthetic RNA sequences to target proteins

**DOI:** 10.1101/2023.07.11.548246

**Authors:** Furkan Ozden, Sina Barazandeh, Dogus Akboga, Sobhan Shokoueian Tabrizi, Urartu Ozgur Safak Seker, A. Ercument Cicek

## Abstract

RNA - protein binding plays an important role in regulating protein activity by affecting localization and stability. While proteins are usually targeted via small molecules or other proteins, easy-to-design and synthesize small RNAs are a rather unexplored and promising venue. The problem is the lack of methods to generate RNA molecules that have the potential to bind to certain proteins. Here, we propose a method based on generative adversarial networks (GAN) that learn to generate short RNA sequences with natural RNA-like properties such as secondary structure and free energy. Using an optimization technique, we fine-tune these sequences to have them bind to a target protein. We use RNA-protein binding prediction models from the literature to guide the model. We show that even if there is no available guide model trained specifically for the target protein, we can use models trained for similar proteins, such as proteins from the same family, to successfully generate a binding RNA molecule to the target protein. Using this approach, we generated piRNAs that are tailored to bind to SOX2 protein using models trained for its relative (SOX10, SOX14, and SOX8) and experimentally validated *in vitro* that the top-2 molecules we generated specifically bind to SOX2.

## 1 Introduction

RNA vaccines came under the spotlight with the promise to end the COVID-19 pandemic as a modular, effective, and secure option for immunization. While this success has proven the effectiveness of RNA molecules as an active means of disease intervention rather than being a passive drug target [24], RNA-based therapeutics are in their infancy [6]. The current therapeutic landscape heavily relies on small molecule drugs [12] which target ∼4% of all human proteins [48] and only ∼25% of deemed druggable proteins [46]. Utilizing RNA to target proteins has a big potential to bridge this gap.

RNA binding proteins (RBP) are abundant in the human body, and they predominantly interact with non-coding RNAs (ncRNAs). Consequently, diseases associated with RBPs exhibit distinct phenotypes based on the specific types of RNA to which these RBPs bind [19]. This specificity is an important aspect in minimizing off-target effects. Furthermore, the capacity for rapid modifications in RNA sequences presents opportunities for personalized treatment approaches and allows for adaptation to evolving pathogens. These advantages further emphasize the potential of RNA-based drugs in the realm of therapeutic interventions. Here, we focus specifically on the piwi-protein-binding family of RNAs, also known as piRNAs, which have shown promise as novel biomarkers for early diagnosis of breast cancer [32] and as therapeutic targets for personalized medicine [53] but have not been used as drugs to the best of our knowledge. They have relatively short sequences (24 to 32 bp), which makes them easier to generate synthetically and to specialize to target certain proteins.

Various techniques have been proposed for engineering and *de novo* RNA sequences, including CRISPR-based methods, structural designs, and fully computational methods. The conventional approach involves tackling the inverse folding problem, where a target secondary structure is provided, and a corresponding sequence is generated. Several algorithms have been employed to design RNA sequences using this method, including both target-specific and general methods [31] [11] [16] [47] [52]. RNAifold and MioRNAifold use a constraint programming-based approach to search for the RNA with the Minimum Free Energy (MFE) leading to the target structure [18] [41]. Other studies use evolutionary algorithms along with constraint-based approaches for designing the target sequences [15] [40]. Similarly, there are models that use a target tertiary structure of the RNA for obtaining the original sequence [51]. On an orthogonal track, a CRISPR-based method is used for designing guide RNAs (gRNA) [36]. It involves editing the RNA sequence using CRISPR and applying various algorithms and rules to select the best versions. A fully in-silico approach is to use a 3D motif library drawn from all unique, publicly deposited crystallographic RNA structures and uses an efficient algorithm to discover combinations of these motifs and helices that solve the RNA motif path-finding problem [54]. When this 3D structural design is completed, they are filled in with sequences that best match the target secondary structure and minimize alternative secondary structures. Although these studies have been successful in generating or modifying RNA sequences to obtain a desired target, the utilized approaches are not able to span the entire space of piRNA sequences due to the limitations of their methodologies. Additionally, these models typically necessitate either an input RNA sequence for a mutation-based optimization or a provided 2D or 3D structure to facilitate sequence design.

More recent approaches benefit from deep learning for designing biological sequences, including DNA [7] and RNA sequences [26] [10] [35] [2]. Huang et al. use a graph neural network for imputation of the RNA sequences [26] as a way to generate the missing sequences obtained from the RNA sequencing devices. Chuai et al. combine the use of a deep learning model with the CRISPR method for designing the RNA sequences [10]. Similar to some older studies, this approach requires a starting RNA sequence and is not capable of generating truly novel RNA sequences. Furthermore, these models do not offer additional control over the functional attributes of the designed sequences, such as their capacity to bind to specific proteins.

## 2 Results

### 2.1 Overview of RNAGEN

The RNAGEN model is a deep generative adversarial network (GAN) that learns to generate piRNA sequences with similar characteristics to the natural ones. This model is a novel version of the WGAN-GP architecture [23] for one-hot encoded RNA sequences. The model provides improved training over the original Convolutional GAN (DCGAN) models [23] [44] and is less prone to overfitting than the original WGAN architecture [4].

Figure 1A shows the overview of the WGAN architecture. The input is a noise vector. The generator and the critic are convolutional neural networks trained together. The generator upsamples the input vector using transpose-convolutions to generate piRNA sequences, whereas the critic uses convolutions and dense layers to score the actual and generated samples. The optimization pipeline is shown in Figure 1B where we select three proteins of the same family that are closest to the target protein in the ProtTrans [14] space and use the distance as weight and the DeepBind [3] predictions as scores, the input vector is learned to generate piRNA sequences with higher binding affinity to the relative proteins and therefore the target protein itself.

**Fig. 1.**
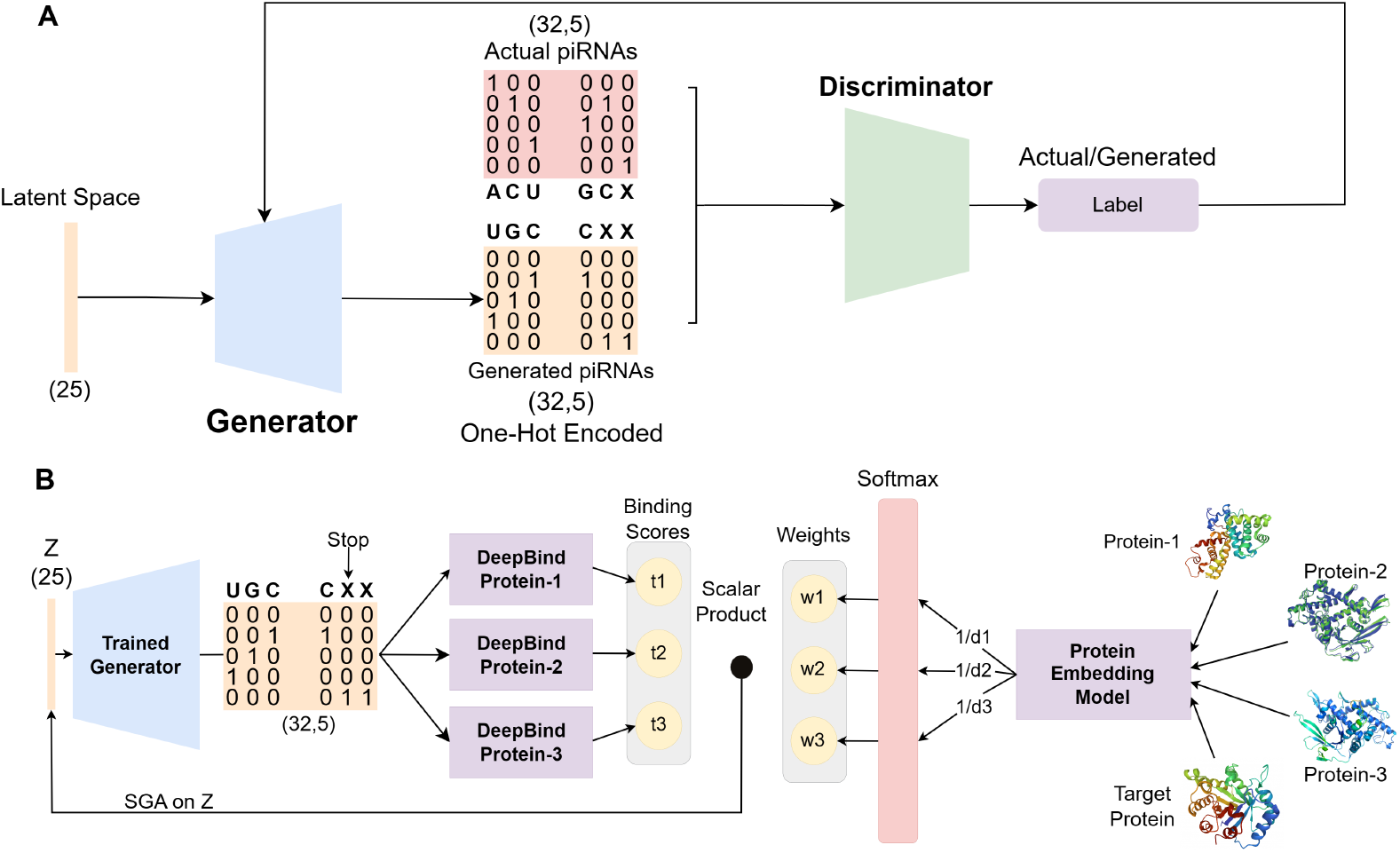
Training and Optimization pipeline. **A** shows the training of the GAN model. The GAN consists of a generator and a critic. The one-hot encoded natural sequences are used to train the model, and the generator also outputs padded one-hot sequences. The sequence is derived from the one-hot encoding, and the conversion stops with the first padding alphabet. **B** is the optimization pipeline, where the generator G is trained on real piRNA sequences. We utilized the ProtTrans model [14] to find proteins similar to our target protein for which pre-trained DeepBind [3] models exist. We demonstrate in Supplementary Figure 12 that the embeddings of homologous proteins extracted from ProtTrans are closer to the reduced dimension space. This indicates that this embedding model is suitable for selecting proxy proteins. RNAGEN learns latent vectors *z* that lead to the generation of optimized piRNA sequences with improved binding scores to the target protein. Our model uses inverse distances (*d*) of relative proteins to the target protein as a weighting tool by which it adjusts the relative improvements to the latent vector *z*. We multiply these weights with the respective binding score predictions from the pre-trained DeepBind models. The result of the scalar product is then treated as a hint score whereby providing a differentiable score function for *z* to optimize for.

### 2.2 Generated sequences maintain their distance to the natural piRNAs

The WGAN architecture described above (and also in detail in Section 4) learns to generate piRNA sequences that resemble the natural piRNA sequences. We compared the distances between synthetic and natural sequences. Due to the variable length of the piRNA sequences, we selected the Levenshtein distance as the distance metric. Levenshtein distance is the minimum number of single-character edits (insertions, deletions, or substitutions) required to change one word into the other [34]. We also trained an AutoEncoder on the same data to compare the performances of these two models and show that the GAN-based model is superior. To evaluate the distance between the set of RNAGEN-generated and AutoEncoder-generated sequences in relation to the natural dataset, we found the distance of each sequence to the closest sequence in the natural samples in terms of Levenshtein distance. Then, we employ these sets of distances to represent the distribution of distances within these sets. To enable a comparison with the natural dataset, we performed a similar analysis on the natural samples by finding the smallest non-zero distance between each natural sample and the entire set of natural piRNA sequences. The distributions shown in Figure 2A demonstrate that the generated sequences exhibit a nearly identical distribution of distances compared to the natural sequences, whereas AutoEncoder-generated samples displayed a distinct distribution. Furthermore, the average distance of the AutoEncoder-generated sequences to the natural sequences is higher than the generated ones, indicating that the generative model has effectively learned the contextual characteristics of the sequences. We used the Mann-Whitney *U* -test here, which is a non-parametric statistical test used to compare differences between two independent sets of samples and is usually preferred over the *t*-test when the data is following non-normal distributions [39].

**Fig. 2.**
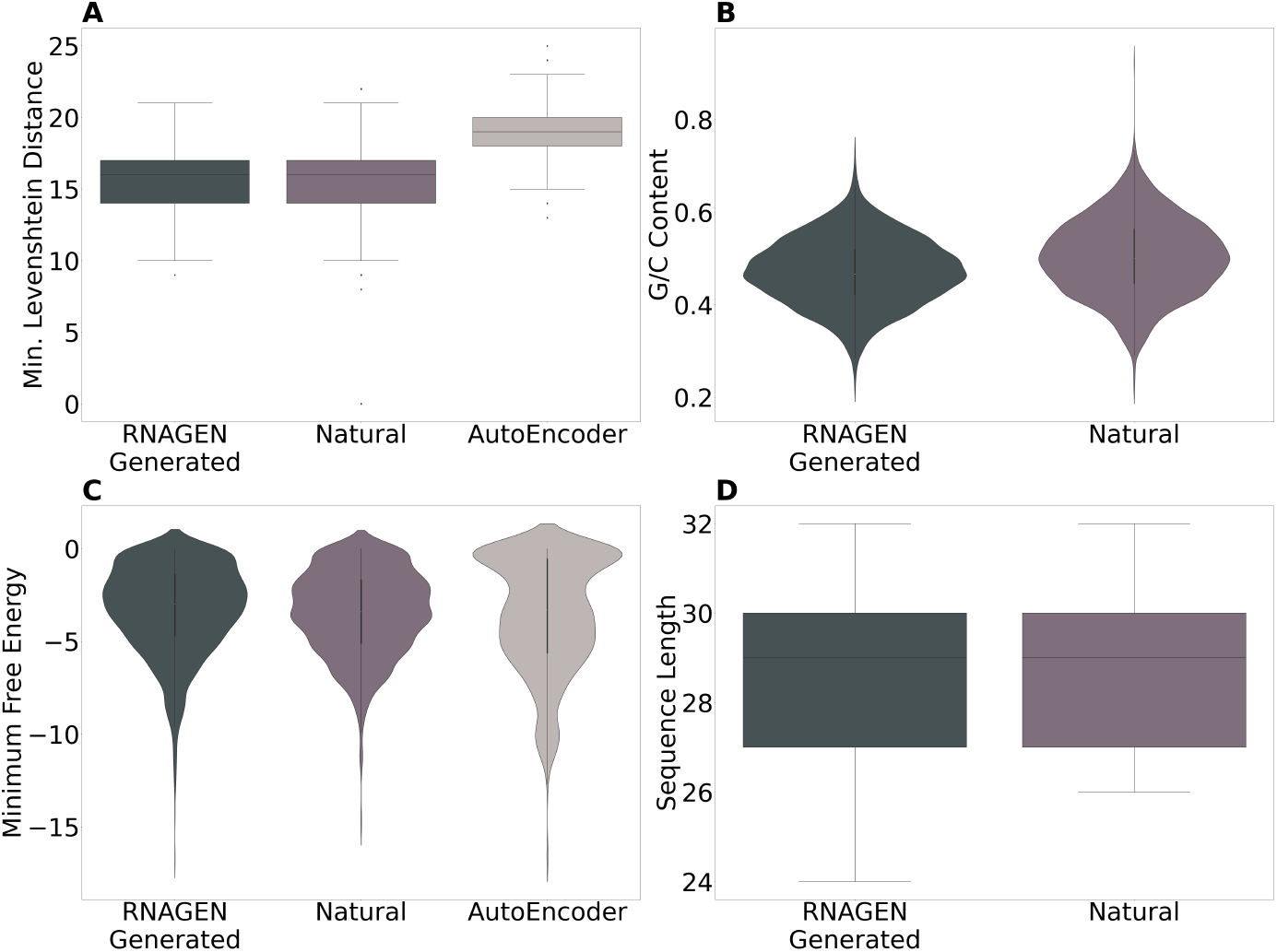
Minimum Free Energy distributions. **A** shows the distribution of the MFE in the RNAGEN-generated, AutoEncoder-generated, and natural piRNA sequences. The AutoEncoder-generated sequences are generated based on the average GC content of the reference genome. The distribution of the MFE in 2048 AutoEncoder-generated sequences is skewed (Mann-Whitney *U* -test: *p* < 0.01) while the 50,379 natural and 19,200 RNAGEN-generated sequences are considerably similar in distribution (Mann-Whitney *U* -test: *p* > 0.01). **B** The distributions of the length of the generated and natural sequences are shown to be similar, with almost identical averages of 28.6 and 28.5, respectively. The length of the sequences affects the MFE, and the results show that the generative model learns to maintain the length as well as the MFE. **A** The Levenshtein distance of the RNAGEN-generated and AutoEncoder-generated sequences to the closest natural sequence is measured, and the distributions of the distances are shown here. We expected the sequences not to be very similar or very distant from the natural set of sequences. To measure this for the actual sequences, the distance to the closest member with a non-zero distance is considered. We use the Mann-Whitney *U* -test for statistical tests, and as shown here, the distance is maintained (Mann-Whitney *U* -test: *p* = 0.18). The *p* value for the AutoEncoder-generated sequences was smaller than 0.01. The size of the AutoEncoder-generated and generated sequences are both 2048, and all the 50,397 natural sequences are used here. **D** The GC content of the 50,397 natural and 2048 RNAGEN-generated sequences are shown with similar mean values. The range of the GC content of the generated one remains in the range of natural sequences while the statistical test does not show similar distributions (Mann-Whitney *U* -test: *p* < 0.01).

### 2.3 GC Content of the natural and generated sequences are similar

We investigated the GC content of the RNAGEN-generated sequences. We calculated the GC content of 2048 generated sequences and compared the distribution to that of 2048 natural RNA sequences. The average length of both sets of sequences was 28 bp. The results in Figure 2B show that the RNAGEN-generated sequences have similar GC content compared to the natural piRNAs. While the average GC content of the entire genome was 59.1%, the GC content of the natural sequences averaged just above 50%. Although the Mann-Whitney *U* -test showed a significant difference (*p* < 0.01) between the distributions of GC content in the natural and generated sequences, the average GC content was similar, and the GC content of the RNAGEN-generated sequences fell within the range observed in the natural ones.

### 2.4 Distribution of the minimum free energy of the generated is similar to that of natural ones

We investigated the MFE of the generated piRNA sequences. MFE of an RNA sequence is the energy of the secondary structure that contributes a minimum of free energy [55]. We used the ViennaRNA package [37] to measure the MFE of the RNA sequences, and the measurements using the Nupack package [17] also yielded similar results. In Figure 2C, we compared the MFE distributions of 19200 generated sequences and the entire set of natural piRNA sequences used for training and validation. The difference in distributions was statistically insignificant (Mann-Whitney *U* -test: *p* > 0.01). Also, we generated a matching number of RNA sequences using the AutoEncoder and compared their MFE distribution with natural and generated sequences. The MFE distribution of the AutoEncoder-generated sequences was substantially different from the counterparts (Mann-Whitney *U* -test: *p* < 0.01). This shows that RNAGAN is able to mimic natural piRNAs in terms of their minimum free energy.

### 2.5 Electrophoretic Mobility Shift Assay (EMSA) confirms binding of two of the optimized piRNA sequences with highest binding specificities

We used our proposed architecture to generate piRNA sequences, specifically designed to have high binding scores to human transcription factor SOX2. Figure 3 shows binding scores predicted by the DeepBind model for the SOX4 (see Supplementary Figure 5A and 5B), SOX2 (see Figure 3C and 3D), and RBM45 (see Supplementary Figure 5C and 5D) proteins, of the generated piRNA sequences before and after performing optimization for each of these proteins separately using our model. Although this model is more useful for proteins without DeepBind prediction models available, here we select proteins (SOX2, SOX4, and RBM45) for which DeepBind models exist to validate the optimization results. In the optimization procedure, 64 piRNAs are generated using the trained generator, and then we optimize the generator’s input latent vector using gradient ascent to maximize the predicted binding score to the target proteins. We validate the optimization using *in vitro* experiments on the SOX2 protein. We observe that the optimization procedure yields piRNA sequences that have high predicted binding scores against SOX2. The maximum predicted binding score obtained is 6.09, which demonstrates a well-binding preference as mentioned in [3]. We also use the AlphaFold 3 [1] server to predict the complexes, including the top 2 RNA aptamers and the SOX2 protein, and observe potential interaction between the aptamers and the target SOX2 protein. As shown in Figures 4 and 5, the prediction of the binding section of the protein and the RNA has a pTM confidence of 0.54 and 0.53, respectively, for aptamer 1 and aptamer 2. The pTM confidence level of the predicted structure of the protein itself is above 0.9 for the interacting region in both complexes [1]. The full protein predictions of the same complexes are shown in Supplementary Figures 9 and 10. The two optimized piRNA sequences with maximum predicted binding scores to SOX2 (dubbed RNA aptamer 1 and RNA aptamer 2) were also studied via *in vitro* validation experiments. We used electrophoretic mobility shift assays (EMSAs) [25]. They analyze interactions between proteins and nucleic acids. The assays use polyacrylamide gel electrophoresis (PAGE) under non-denaturing conditions and buffers at or near neutral pH. The migration of free and bound biomolecules in an electric field is measured to determine the affinity and specificity of the interaction [45, 49, 50]. Here, we used an agarose gel electrophoresis technology that has been developed to allow for short run times while simultaneously yielding high band resolution, avoiding short-RNA aptamers from degrading, and detecting RNA-bound protein from free RNA with a nucleic acid dye, which is safer than conventional radioactive labeling [43]. *In vitro* binding reactions were visualized using agarose gel-based rapid EMSA, which was performed to verify the binding between obtained RNA aptamers and SOX2 protein (Figure 6). The arrow in Figure 6, indicates the *in vitro* binding of piRNA sequences and purified SOX2 protein, demonstrating saturation as the RNA concentration increases. Unbound RNA, being molecularly smaller than the RNA-protein complex, moves faster in the agarose gel with applied voltage. The gradient of this movement is marked with a bracket. The bands indicated by the arrow suggest that piRNAs, RNA aptamer 1 and RNA aptamer 2, bind to purified SOX2.

**Fig. 3.**
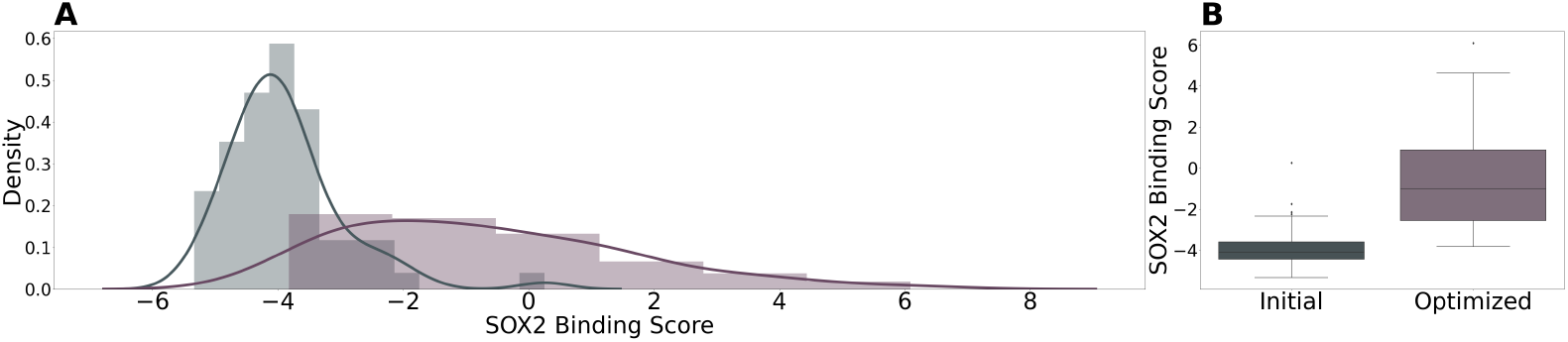
Predicted binding scores before and after optimization. **A**,**B** Show the initial and optimized predicted binding scores of the generated piRNA sequences to the SOX2 protein using SOX10, SOX14, and SOX8 proteins as proxy proteins for optimization. In this case, not only is the maximum predicted binding score an order of magnitude larger, but the highest score is also a considerably high score of 6.09.

**Fig. 4.**
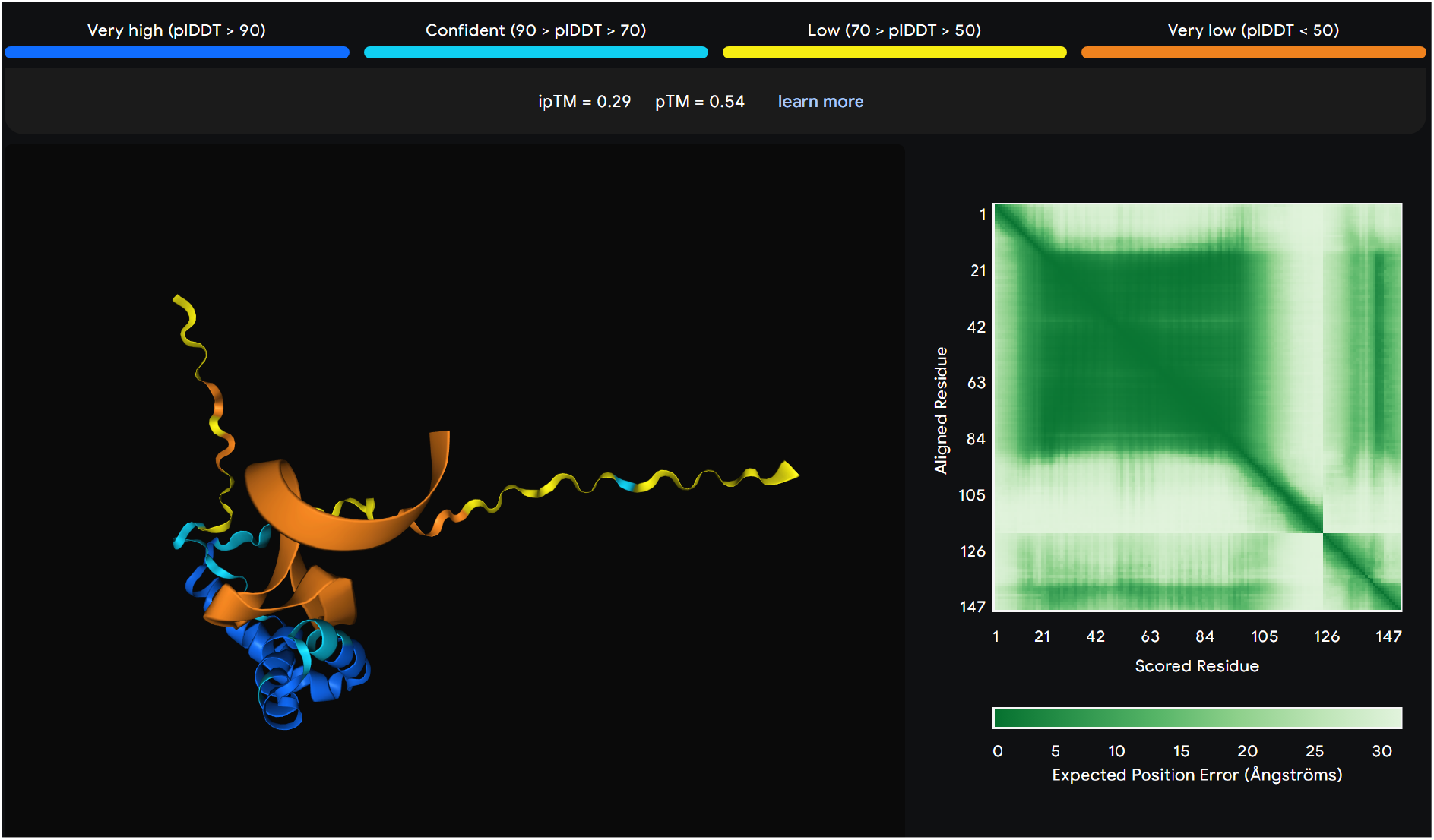
Prediction of the structure of the binding region in a complex including Aptamer 1 and the SOX2 protein using AlphaFold 3. The confidence levels here for the full complex and the selected protein region are 0.54 and above 0.9, respectively.

**Fig. 5.**
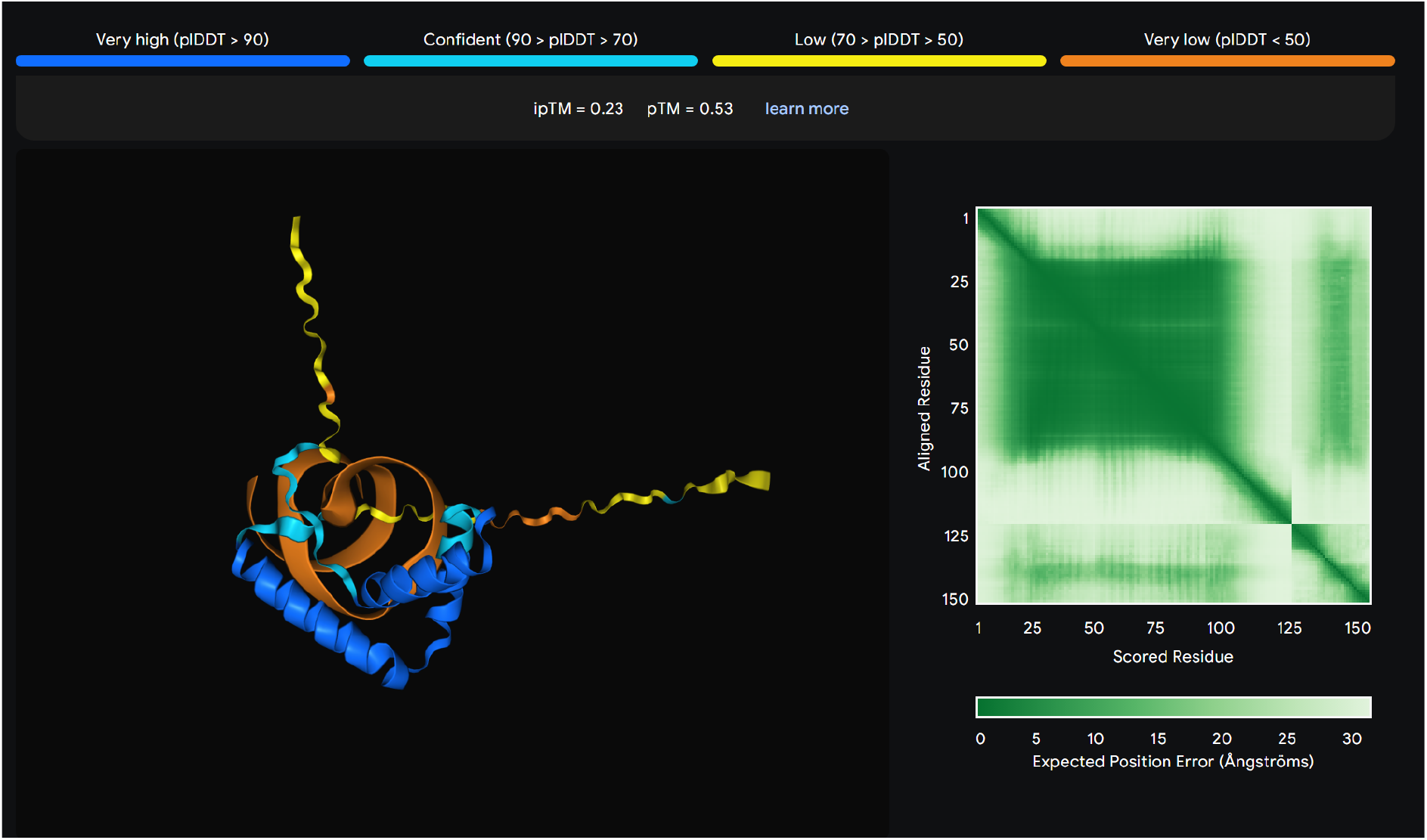
Prediction of the structure of the binding region in a complex including Aptamer 2 and the SOX2 protein using AlphaFold 3. The confidence levels here for the full complex and the selected protein region are 0.53 and above 0.9, respectively.

**Fig. 6.**
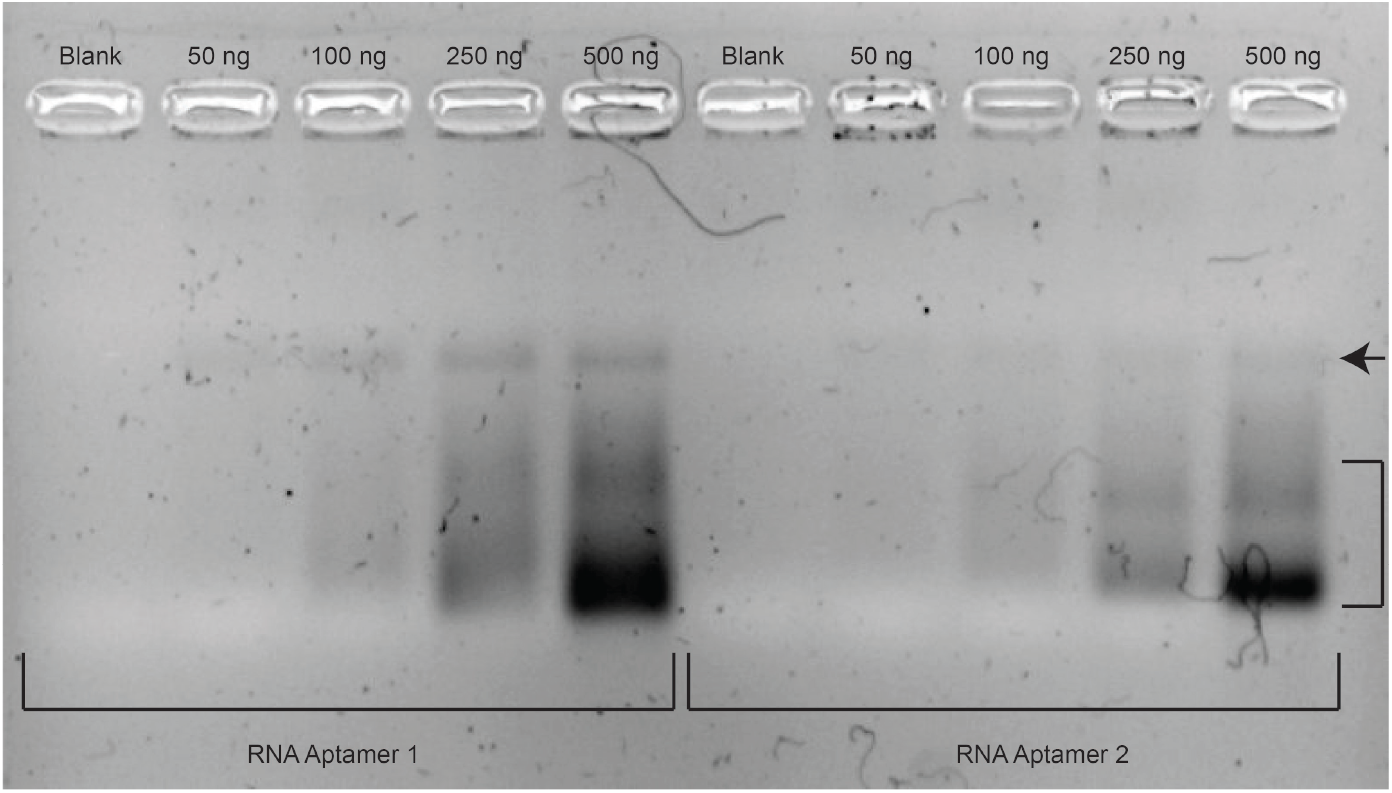
Rapid agarose gel electrophoresis for electrophoretic mobility shift assay. Binding reactions with a set amount of purified SOX2 protein (2.5 *µ*g) and a concentration gradient of RNA aptamer (The amounts of RNA are equivalent to 430 nM - 4.3 *µ*M for RNA aptamer 1 and 460 nM - 4.6 *µ*M for RNA aptamer 2) were loaded onto the gel. The saturation of RNA-protein binding is indicated by an arrow. The shift of RNA aptamer-protein binding to free RNA is indicated by a curly bracket.

To measure the specificity of the binding computationally, we measure the binding score of the top 2 generated piRNAs to all DeepBind models available, which includes more than 270 models. We observe that only a few other proteins have a considerable likelihood of binding, some of which are proteins from the same family as SOX2. See Supplementary Figures 7 and 8 for details.

**Fig. 7.**
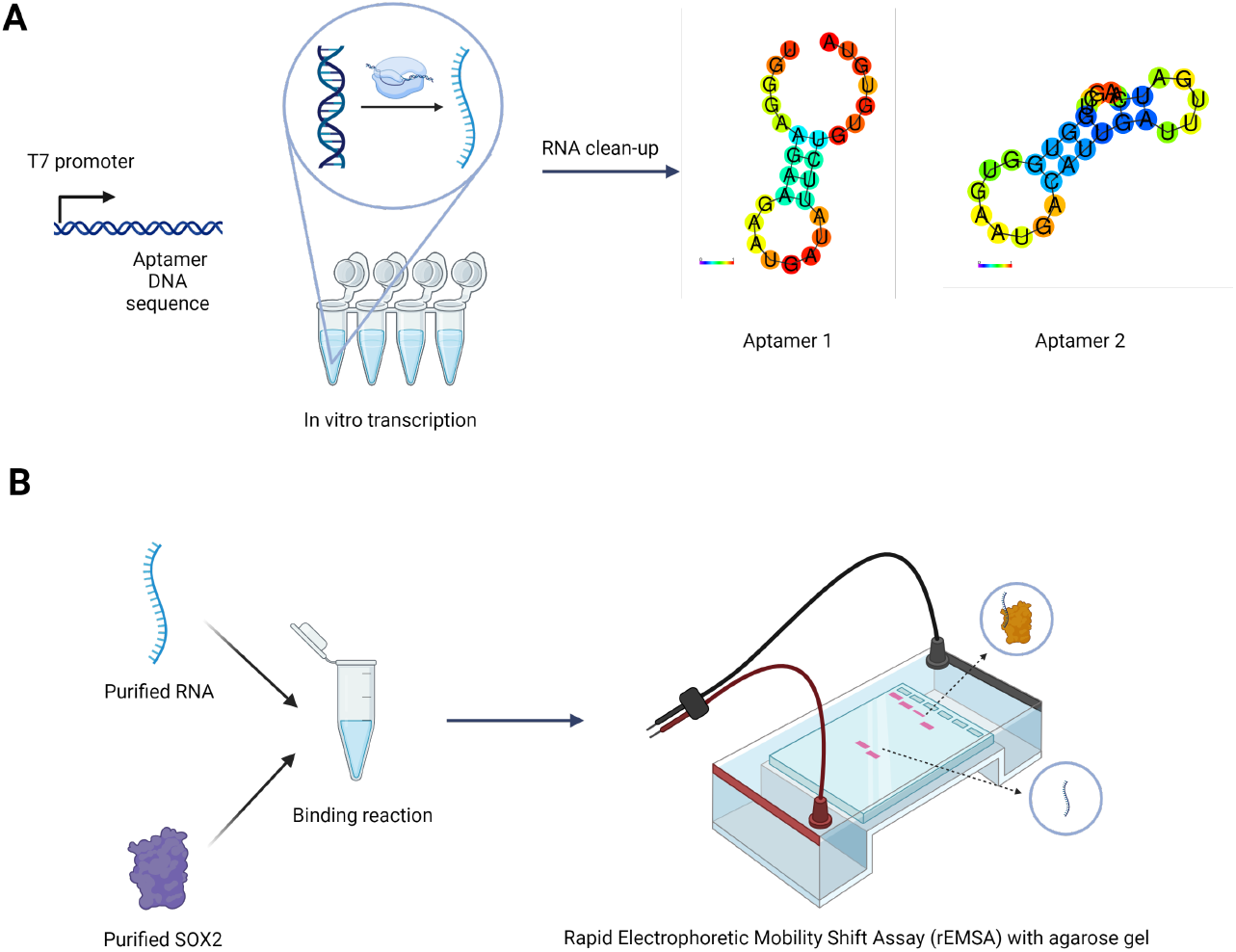
Experimental pipeline. (A) RNA aptamers were produced using in vitro transcription described by [38]. Produced RNAs were purified and eluted into the binding buffer. The secondary structures of RNA aptamers were obtained from RNAfold WebServer [22]. (B) Purified RNA and purified SOX2 proteins are mixed for binding reaction and run on agarose gel for rapid Electrophoretic Mobility Shift Assay (rEMSA). Bands were visualized with SYBR Safe dye. The image was created with BioRender.com.

## 3 Discussion

We introduce RNAGEN, the first framework for generating optimized piRNA sequences with desired binding properties using indirect optimization. Our approach enables us to generate sequences with similar attributes to the natural piRNA samples without the need for exploring intractable search space of RNA sequences that have lengths of up to 32 nucleotides. Our model can also generate sequences of variable lengths, further increasing the possible space of generated sequences.

RNAGEN is a robust framework for synthetic piRNA engineering and is applicable to numerous applications for genetic circuit design. The model’s components consist of a GAN adopted to be trained on the human natural 5’ piRNA dataset and state-of-the-art protein-binding prediction models as well as protein embedding models. Our model can potentially optimize sequences for any objective function as long as a differentiable prediction model exists to predict the target metric for a piRNA sequence. Similarly, the model can work with classification models to generate sequences of a particular class if the prediction model is replaced with such a classifier.

Our model does not rely on the availability of predictions for the binding scores of the generated piRNA sequences and can optimize piRNAs for the target protein using proteins similar to the target protein in the ProtTrans [14] space. For this purpose, proteins of the same family for which RNA-protein binding prediction models exist are selected. By increasing the weighted average binding score for these three proteins, the average predicted binding score of the generated piRNA sequences to the target protein is expected to increase.

While RNAGEN is capable of generating and optimizing sequences with favorable binding scores, the optimization process does not consistently yield sufficiently high predicted binding scores. One potential explanation for this outcome is the constraint imposed by the predictor model (DeepBind), which is not perfectly accurate. Moreover, the selection of relative proteins for the optimization process is based on their similarity in the ProtTrans embedding space, which may not align with their similarity in the DeepBind models’ space. Another limitation of RNAGEN is its reliance on available DeepBind models to select proteins similar to the target protein, disregarding the absence of DeepBind models for certain proteins. Consequently, the top three selected proteins might lack the necessary similarity to facilitate meaningful optimization. Thus, when using proteins selected from the ProtTrans embedding space, the enhancement in the predicted binding score for some proteins may be relatively less significant, as seen in Supplementary Figure 5A and 5B for the SOX4 protein. Additionally, in rare cases, optimization may lead to decreased predicted binding scores. However, given the flexibility of the optimization steps, if no improvement is achieved after multiple iterations, the best initial predicted binding score can be utilized.

We expect to see wide use of data-driven sequences generated using deep learning methods in the near future. Generating RNA utilizing this approach is practical and does not require massive computational resources. While we focus on piRNA, it is straightforward to generalize the framework to other families of RNA sequences or specific regulatory elements. It is also possible to use other generative models, such as transformers or diffusion models, for more complicated tasks or families of longer sequences. Generating piRNA sequences with specific binding targets enables researchers to conduct experiments in a much shorter time, which can be particularly beneficial in RNA-based drug design.

Our *in vitro* experiment, rapid-EMSA, indicates a promising specificity of the top-2 generated piRNA sequences towards the target, SOX2 protein. Although not quantified, the observed increasing band intensity at protein-RNA binding and saturation of this binding provides preliminary evidence of binding specificity (Figure 6). This result is further supported by the lack of a saturation gradient when the top-2 generated piRNA sequences were tested against Bovine Serum Albumin (BSA), a protein not targeted by our piRNAs (Figure S5). However, to further substantiate these findings, it is essential to quantify the binding rate and the mode of the affinity of these piRNAs to their target protein.

A combination of techniques can be used to elucidate the binding dynamics of our generated piRNAs. Surface Plasmon Resonance (SPR) can monitor binding kinetics in real-time, providing a practical understanding of the interaction rates [28]. To further comprehend the in vitro binding affinity of piRNAs to their target protein, an RNA pull-down technique that allows for the isolation and subsequent identification of proteins that specifically bind to the piRNAs can be employed [9]. Lastly, it is essential to utilize a combination of techniques for the reliable characterization of RNA-target interactions. For instance, the use of isothermal titration calorimetry (ITC), mass spectrometry (MS), and nuclear magnetic resonance (NMR) could provide comprehensive insights into again the binding affinity, selectivity, and mechanism of the generated piRNAs toward their target protein [8]. These methods could offer a more applicative view of the binding dynamics of our piRNAs.

RNA molecules generated with RNAGEN that display specific protein targeting could be utilized as aptamers in RNA-based therapeutics or as sensory elements in molecular sensors for tracking diseases and identifying cellular phases [13, 42]. For instance, these RNA molecules generated with this technique can be enhanced by increasing the stability of their secondary structures in their active conformation and be delivered in synthetic mRNA devices to distinguish cell types [29]. Our top two piRNAs targeting SOX2 protein could be utilized again in synthetic genetic circuits to track SOX2 trajectories and monitor reprogramming efficiency during induced pluripotent stem cell (iPSC) generation [27]. Furthermore, the generated RNA molecules could be utilized to control gene expression by targeting transcription factors in metabolic engineering [30].

## 4 Methods

### 4.1 Data set

Our proposed method needs an RNA sequence data set to train the GAN. Even though any computationally suitable RNA type can be modeled, in this work, we focused on piRNAs due to their shorter length, as mentioned before. This characteristic makes it easier for the GAN to learn the underlying sequence patterns effectively. To train the generator model, we used the DASHR2-GEO piRNA database [33]. The dataset contains 50, 397 natural piRNA sequences of lengths varying between 26 and 32 bps. The distribution of the length of these sequences is shown in Figure 2B and compared to those of synthetic piRNA sequences. We also tried to generate and optimize sequences using simulated annealing and genetics algorithms; however, these methods do not learn anything from the natural sequences and rely on mutations and crossover operations over random sequences. In addition, these approaches do not guarantee to reach the objective characteristic in any certain number of iterations.

### 4.2 Experimental Setup

We encode the data using one-hot encoding, and the dimension of each RNA is (32, 5), where 32 is the maximum size, as mentioned, and 5 is the length of the one-hot vector. The batch size during the training and optimization is 64, and we randomly divide the dataset into training and validation sets where the validation set consists of 12.5% of the sequences. The RNA alphabet in one-hot encoded as follows: *A*:(1,0,0,0,0), *C* :(0,1,0,0,0), *G* :(0,0,1,0,0), *U* :(0,0,0,1,0), *X* :(0,0,0,0,1) where *X* is the padding character used when the sequence is shorter than 32 bps. This padding is necessary because the input of the discriminator (critic) as a convolutional neural network has to be in fixed dimensions, and the sequences naturally vary in length. For details regarding hardware settings, see Supplementary Notes. Details regarding hyperparameter optimization can be found in Supplementary Notes 4. The code is available at https://github.com/ciceklab/RNAGEN.

### 4.3 Problem Formulation

In this work, we assume that only the sequence information of the target protein is known, and no other computational tool is available to assess the binding score of any given RNA sequence to the target protein. If there is such a computational tool available, the availability of such a computational tool requires more information than the sequence information alone, and then the optimization procedure is more straightforward. Let *P*_*t*_ be the sequence information of the target protein for which we want to design a piRNA to have a high binding preference to it. We assume that a cohort of binding score assessment models for well-known proteins (e.g., DeepBind models) are available. Let the family of assessment functions be {*F*_*i*_(*R*)} where *R* is a variable representing any possible input RNA sequence and the corresponding protein sequence information for the function *F*_*i*_(*R*) be *P*_*i*_. Since {*F*_*i*_(*R*)} does not contain *F*_*t*_(*R*) (i.e., the assessment function for target protein), we imagine a latent function *RB*_*t*_(*R*) which returns a ground truth real binding score of any input RNA sequence to the target protein. Then, our goal is to find an optimal *R* = *R*^∗^ such that arg max *RB*_*t*_(*R*) = *R*^∗^. This is without knowing any information on the latent function *RB*_*t*_(*R*) while using *P*_*t*_, {*F*_*i*_(*R*)} and {*P*_*i*_}.

### 4.4 Model

The architecture shown in Figure 1 is composed of (i) a trained generator, (ii) the required number of trained property assessment models, and (iii) a trained protein embedding model. The model requires two optimization procedures. The first required optimization procedure where a generative model (see Figure 1A) consisting of a generator G and a critic C is trained. In this architecture, a GAN is used to capture the latent piRNA distribution and limit the optimization space of the random noise *Z* (i.e., the second required optimization procedure shown in Figure 1B) with the realistic piRNA subspace. The employed approach provides a realistic optimization on the latent noise *Z* since each time it passes through the generator, the resulting generated piRNA sequence must belong to the subspace spanned by the trained generator. The purpose of the overall architecture is to find an optimal *Z* = *Z*^∗^ (i.e., latent noise for the generator), such that *G*(*Z*^∗^) outputs the desired RNA sequence with maximum binding preference to the targeted protein, where *G* is the generator function.

#### Generative Model Architecture

In this work, we utilize Generative Adversarial Networks (GAN) [21] as the generative model. The input of the GAN is a random noise sample *z*, and we train the GAN to generate realistic RNA sequences. The GAN learns to map the noise to the sequence space. To optimize the sequences, we use gradients to update the input noise only. In this optimization, the generalized model is preserved, and the outputs are optimized based on the desired objective.

The layers of the deep learning model are modified to fit this task based on the dimensions of the data, with 32 nucleotides, each represented by a vector of length 5. The generator consists of a dense layer to increase the dimension of the input noise, following five residual convolution blocks, all using the same number of channels. Finally, the output of the last residual block is fed to a convolution layer with five output channels following a Softmax layer, the output of which is the one-hot encoded sequence (see Table S3). The input is a random vector of length 25, sampled from a normal distribution with a mean of zero and a standard deviation of 1.0. Technically called critic, the discriminator of this model is a convolutional neural network with five residual blocks following a dense layer with one output (see Table S4). Unlike the original DCGAN model [44], there is no activation function at the output of the dense layer of the critic, as the output is an unbounded score and not a probability [4]. The number of iterations of the critic per generator iteration is five here, meaning that the critic is updated five times more than the generator.

As mentioned above, the generative model consists of a generator *G* and a critic *C*. The generator gets a random noise as input and generates random samples, and the weights of the network are initialized with a normal distribution range of -0.1 and 0.1. The model’s loss function, as shown in Equation 1, is to maximize the distance between the critic’s scores for the actual sequences and the generated ones. The first part of the loss function belongs to the original WGAN [4], and the second part includes the gradient penalty and its coefficient *λ*. Here *x* and 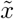 are the actual and generated samples, where we sample 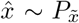 uniformly along straight lines between pairs of points sampled from the data distribution ℙ*r* and the generator distribution ℙ*g* for the computation of the gradient penalty. The gradient penalty improves the training of the GAN by enforcing the Lipschitz constraint as the model proposes [23], and this allows training for a higher number of iterations. Here, we train our model for 200,000 generator iterations.

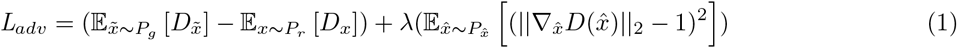

#### Protein Embedding Model

The proposed architecture needs a similarity metric between proteins in order to make use of the similarities between target and related proteins and make inferences about the binding preferences of generated RNA sequences to the targeted protein. In this work, we use the protein embedding model proposed in [5] to extract the 100*d* vector representations of the protein sequences. Using these vector encodings, we calculate the pair-wise Euclidean distances of all proteins, for which a binding assessment model is available, with the targeted protein. The protein embedding model is a deterministic function *f*_*e*_(*P*), where *P* is the sequence information of the RNA. It uses each 3−mer in the protein amino acid sequence and, using a look-up table (i.e., embedding vectors), it calculates the overall protein embedding by aggregating the embedding vectors for each 3−mer. We summarize the function of the protein embedding model via the following relation *X* = *f*_*e*_(*P*) where *X* and *P* denote the protein embedding vector and corresponding protein amino acid sequence, respectively.

#### Target Feature Assessment Models

In order to guide the generative optimization process, our model needs guidance by which it learns to generate piRNA sequences that have a high binding preference to the targeted protein. To serve this *guidance* purpose, our model uses what we call target feature assessment models. Although, in this work, we are using protein binding as our choice of the target feature, it is possible to use any computationally or experimentally assessable target feature to optimize for (e.g., translation rate, half-life).

In this work, we make use of pre-trained DeepBind models. DeepBind models use CNN-based deep networks to evaluate the binding score of any RNA sequence to the protein of interest. DeepBind provides us with a vast array of proteins for which there exists a pre-trained assessment model.

We evaluate the binding score of the generated sequence at each iteration of the generative optimization to a set of proteins of interest. We denote the assessment functions with *F*_*i*_(*R*), where *i* enumerates available pre-trained assessment models and *R* denotes the input RNA sequence. The pre-trained assessment models output target feature scores by which the model guides its generative process. We summarize the assessment relation with *y*_*i*_ = *F*_*i*_(*R*), where *y*_*i*_ denotes the assessment score of *R*, the input RNA sequence, w.r.t. assessing function *F*_*i*_.

### 4.5 Training and Optimization

#### Combiner Block

Our model is able to generate piRNA sequences for proteins even when a target feature assessment model is not available for the target protein. The model only needs the amino acid sequence of the protein to be optimized. Using the corresponding embedding of the target protein, it searches the available assessment models database and finds the proteins which have protein embeddings with minimum Euclidean distance to the target proteins embedding vector. Using a pre-defined number, which we introduce as a hyperparameter to the model, of the closest proteins (e.g., three of the closest proteins), our model weighs the assessment scores inversely with the corresponding Euclidean distance to the target protein. That is:

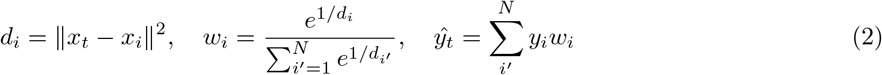

where, *d*_*i*_ denotes the distance of the *i*th closest protein’s embedding to the target protein embedding, *x*_*t*_ denotes the target protein embedding, *N* denotes the number of closest proteins to be used, *w*_*i*_ denotes the corresponding scalar weight for the *i*th closest protein and *y*_*i*_ denotes the assessment score associated with the *i*th closest protein. Hence, the combiner block provides an estimate of the latent function *RB*_*t*_(*R*), which we denote by *ŷ*_*t*_. Notice how *ŷ*_*t*_ = *RB*_*t*_(*R*) when *N* = 1 and the target feature assessment model for the target protein is available. We decide the number of proxy proteins by looking at the distribution of the distances of each protein to its relatives. See Supplementary Figure 11 for more details.

#### Target Feature Optimization

The overall architecture tries to maximize the estimate of the target protein binding score *ŷ*_*t*_ by iteratively updating the input latent noise *Z* with Stochastic Gradient Ascent (SGA). The generation process is described in Algorithm 1 in Supplementary Notes. This algorithm shows the steps for optimizing a single sequence, and we perform this for a batch of sequences and select the best version of each sequence independently.

### 4.6 *In vitro* experiments

Processes of the production and purification of RNA aptamers and SOX2 protein and rEMSA of binding reaction were described in Figure 7.

#### Plasmid design and construction

The human SOX2 gene (RefSeq ID: NM003106.4) nucleotide sequence with N-terminal 6*x* His-tag was chemically synthesized (Genewiz, NJ, USA). With primers containing suitable overhang sequences homologous to the pET22b(+) cloning vector, the SOX2 gene was cloned into the pet22b(+) vector using Gibson assembly [20]. The reaction was transformed into a chemically competent strain of E. coli DH5*α* PRO by heat-shock transformation. Selected colonies were verified by NGS Sequencing (Intergen, Turkey). The SOX2 amino acid sequence is listed in Table S1. All of the primers used in this study are listed in Table S2. The pET22b(+) backbone with SOX2 gene map is shown in Figure S2. Sequencing data is shown in Figure S2. Analysis of NGS data was done using Geneious Prime 9.1.8. software.

#### SOX2 production and purification

The construct was transformed into the E. coli BL21 (DE3) strain for expression. Cells from an overnight culture were diluted to 1:100 in LB medium with the proper antibiotics and incubated until the OD_600_ was between 0.4 and 0.6. The culture was then induced with 1 mM IPTG for 24 hours at 16 °C and 200 rpm. Using a centrifuge, the cell pellet was collected and resuspended in the lysis buffer (20 mM NaH_2_PO_4_, 500 mM NaCl, 20 mM imidazole pH 7.4) supplemented with lysozyme (1mg/mL) and PMSF (1mM). First, five cycles of freeze/thaw were applied. Then, for ten cycles of 15s on/45s off, the suspension was sonicated at 30% power. The samples were centrifuged for 1 hour at 21,500 g, and the supernatant was then filtered through a 0.45 m filter. Using ÄKTA start protein purification system (GE Life Sciences), the filtered lysate was loaded onto the HisTrap nickel column (GE Life Sciences 17524701) that had already been pre-equilibrated. 10-column volumes of lysis buffer (20 mM imidazole) and 5-column volumes of lysis buffer with an imidazole gradient from 50 to 100 mM were used to wash the column. Finally, five volumes of elution buffer were used to elute proteins (20 mM NaH_2_PO_4_, 500 mM NaCl, 500 mM imidazole pH 7.4). Using a HiTrap desalting column (GE Life Sciences, 29048684) in ÄdKTA start protein purification system, purified SOX2 was desalted into 5 mM Tris–HCl buffer (pH 8.0). Protein concentration is calculated via BCA colorimetric assay with BSA standards (Thermo Fisher Scientific, 23225). Protein production was verified using Western Blotting (Figure S3).

#### RNA aptamer production and visualization

RNA aptamers were produced as described in [38]. Aptamer sequences with T7 promoter sequences at 5’-end *in vitro* were transcribed using HiScribe^®^ T7 High Yield RNA Synthesis Kit (NEB, E2040S), according to the kit’s manual. RNA purification was performed using Monarch^®^ RNA Cleanup Kit (NEB, T2040S) according to the kit’s manual. Visualization of the RNA aptamers was done using 2x TBE Agarose Gel Electrophoresis (0.13 M tris (pH 7.6), 45 mM boric acid, 2.5 mM EDTA, 0.02 g/mL agarose, 5x SYBR Safe (Thermo Fisher, S33102)) using RNA loading dye (NEB, B0363S) (Figure S4). Gels were visualized using the Fusion Solo S imaging system (Vilber). Gel images are processed using Fiji/ImageJ software.

#### Aptamer-binding reactions and rapid Electrophoretic Mobility Shift Assay (rEMSA)

Binding reactions and gel preparation were performed as described in [43]. Both protein and RNA were suspended in a binding buffer (5 mM Tris–HCl, pH 8.0). Samples were incubated for 20 min on ice and mixed with a 6x loading buffer. High-percentage agarose gel was prepared using 0.5x TB buffer (45 mM Tris base, 45 mM boric acid, pH 8.2) and agarose (0.03 g/ml), 5x SYBR Safe (Thermo Fisher, S33102). The same binding reaction setup was implemented in testing any nonspecific binding occurring with serum albumin protein (BSA) (SERVA, 11930.01) and observed no shift (Figure S5). Reaction volumes were set to 12 *µ*l. 2.5 *µ*g of protein amounts were used in the binding reactions. Gels were electrophoresed at 200V for 20 minutes. Gels were visualized using the Fusion Solo S imaging system (Vilber). Gel images are processed using Fiji/ImageJ software. Processes of the production and purification of RNA aptamers and SOX2 protein and rEMSA of binding reaction were described in Figure 4. The secondary structures of RNA aptamers were obtained from RNAfold WebServer [22].

## Competing Interests

The authors declare no competing interests.

## Author Contributions

SB and FO designed and implemented the computational model. DA performed the in vitro experiments. SB, FO, DA, SS, UOSS, and AEC wrote the manuscript. UOSS and AEC supervised the study.

## Data Availability

All the data used here are available in public repositories. We downloaded the dataset from the DASHR2-GEO piRNA database [33]. The additional data used in the optimization procedure are available in the public DeepBind repository.

## Code Availability

The source code, including the implementation of the model, the optimization procedure, and the additional scripts used to generate plots, predict binding scores, and find the closest relative proteins, are released at http://github.com/ciceklab/RNAGEN.

## Notes

https://github.com/ciceklab/RNAGEN

### Competing Interest Statement

The authors have declared no competing interest.

### Summary of Updates

We have confirmed the in-silico results using the Alphafold 3 server and have included further analysis of the previous results.

