## Supplemental Material for "RNAGEN: A generative adversarial network-based model to generate synthetic RNA sequences to target proteins"

#### 1 Supplementary Figures

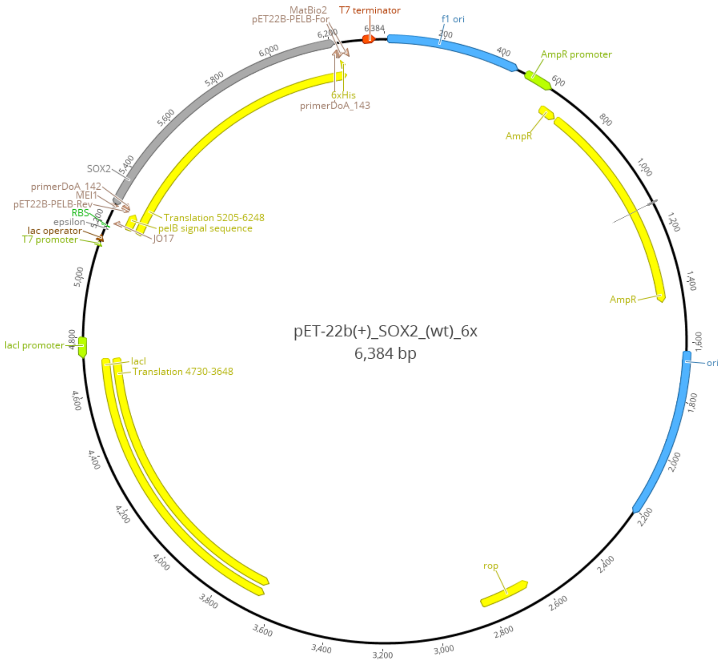


**Fig. 1.** Plasmid map for the production of SOX2.

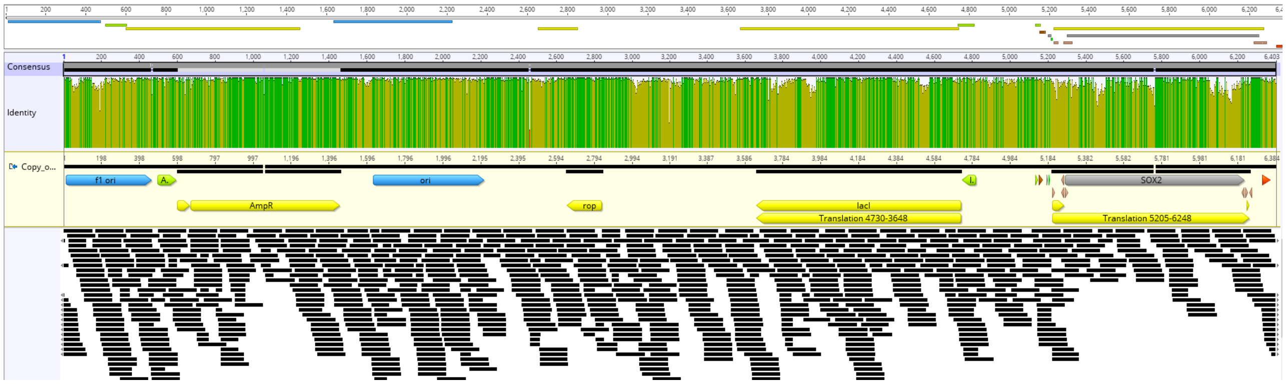


**Fig. 2.** NGS verification of the plasmid used in this study.

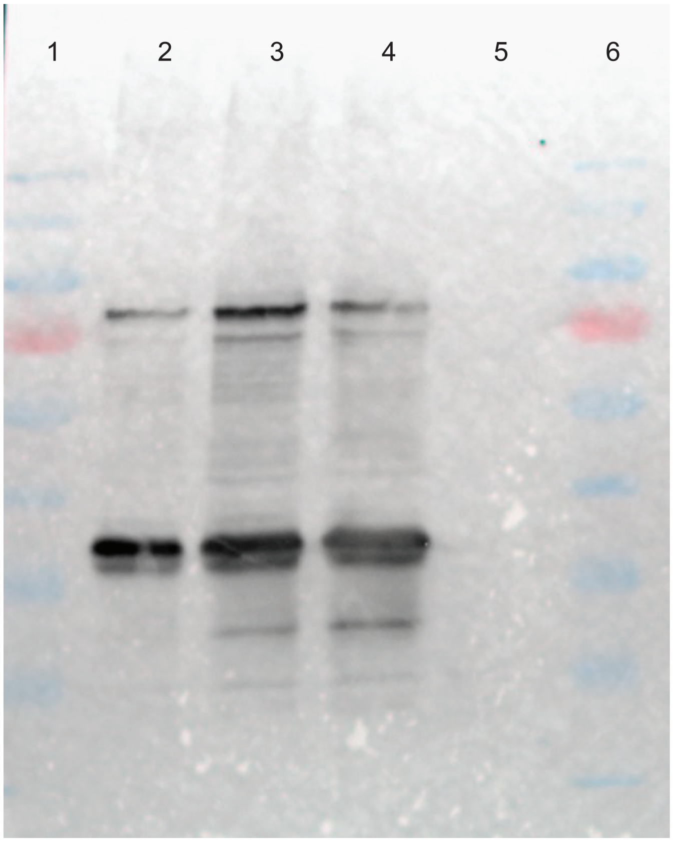


**Fig. 3.** Western Blot verification of SOX2 protein production. Lanes: (1 and 6) PageRuler™ Prestained Protein Ladder, 10 to 180 kDa (Thermo Fisher, 26616), (2-4) SOX2 from BL21 cells, pelB-SOX2-6His – 37.5 kDa, SOX2-6His – 35.3 kDa, (5) BL21 blank as negative control.

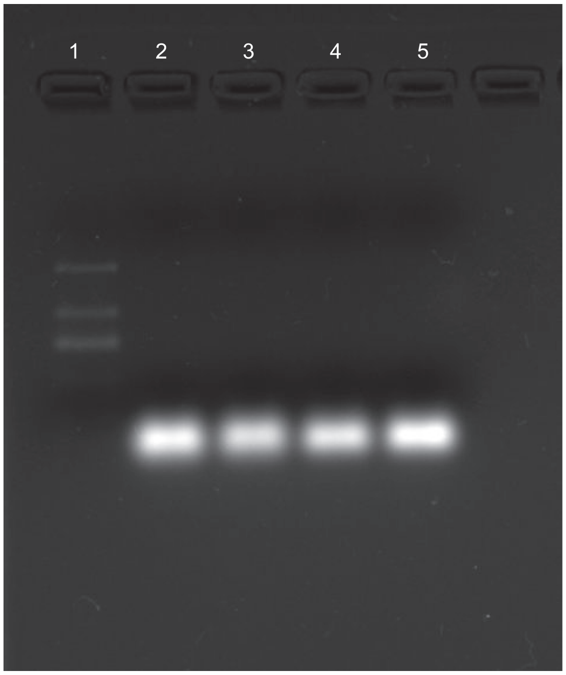


**Fig. 4.** TBE gel electrophoresis verification of RNA aptamer production. Lanes: (1) ssRNA low range ladder (NEB, N0364SVIAL), (2-3) RNA aptamer 1, (4-5) RNA aptamer 2.

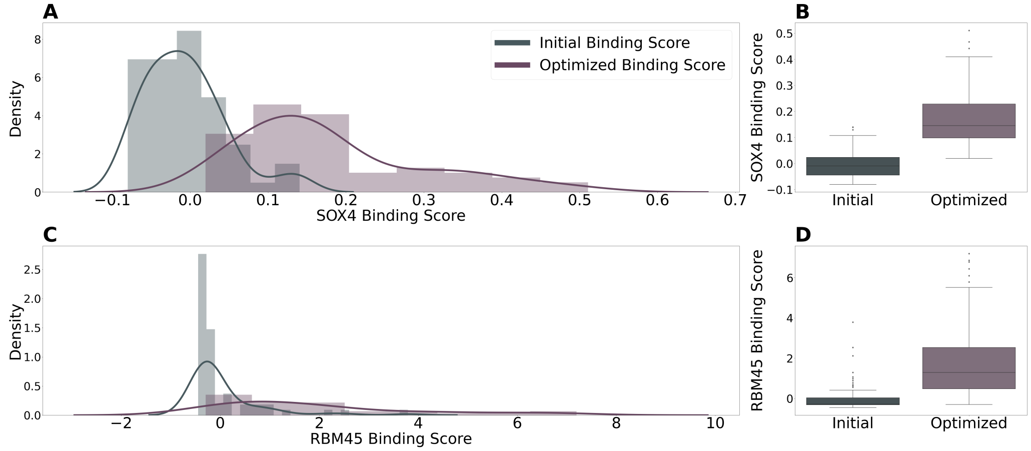


**Fig. 5. Predicted binding scores before and after optimization. A**,**B** Show the distribution of predicted binding scores of the initially generated and optimized piRNA to the SOX4 protein using DeepBind models of the SOX8, SOX7, and SOX10 proteins. As shown in both plots, the predicted binding scores increased substantially, and there was a three-fold increase in the maximum predicted binding score after optimization. **C**, **D** Are the optimization results for the RBM45 protein. Similar to SOX2, the optimization results in high binding scores and reaches up to 7.19 compared to the maximum score of 3.79 for initially generated sequences. The proteins from the RBM family used for optimization are RBM46, RBM41, and RBM4.

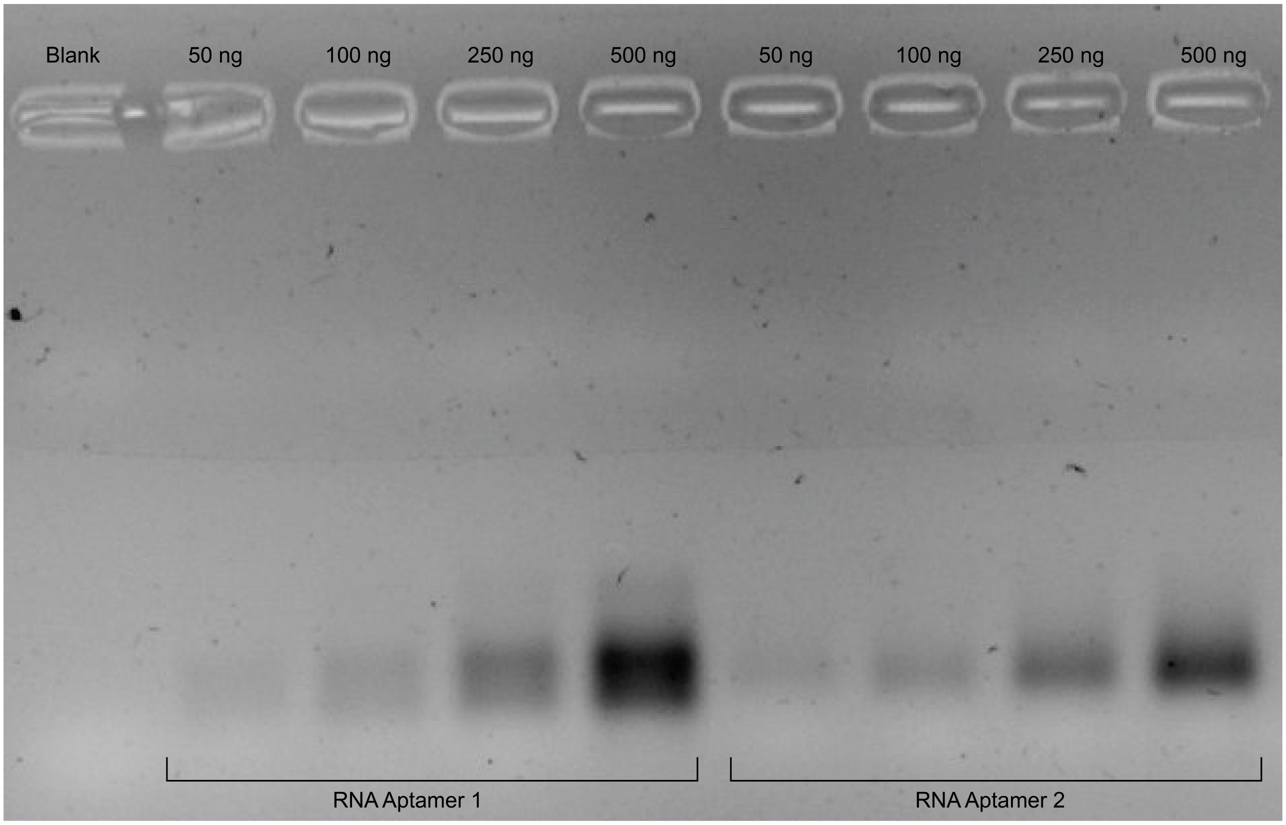


**Fig. 6.** Rapid agarose gel electrophoresis for BSA-aptamer binding control assay. Binding reactions with a set amount of BSA protein (2.5 $\mu$g) and a concentration gradient of RNA aptamer (The amounts of RNA are equivalent to 430 nM - 4.3 $\mu$M for RNA aptamer 1 and 460 nM - 4.6 $\mu$M for RNA aptamer 2) were loaded onto the gel.

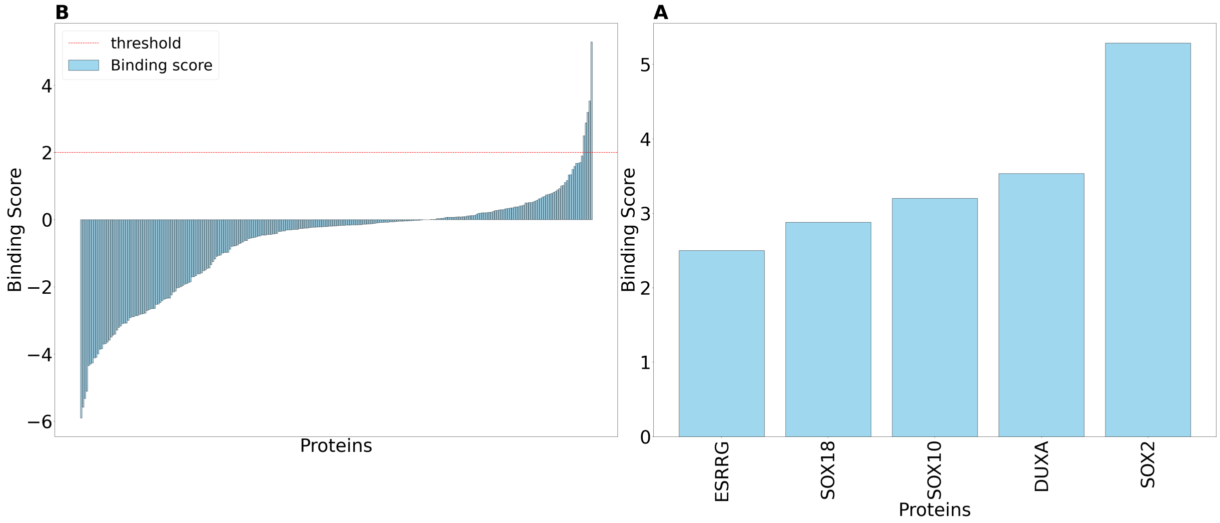


**Fig. 7. Specificity of Binding for Aptamer 1.**

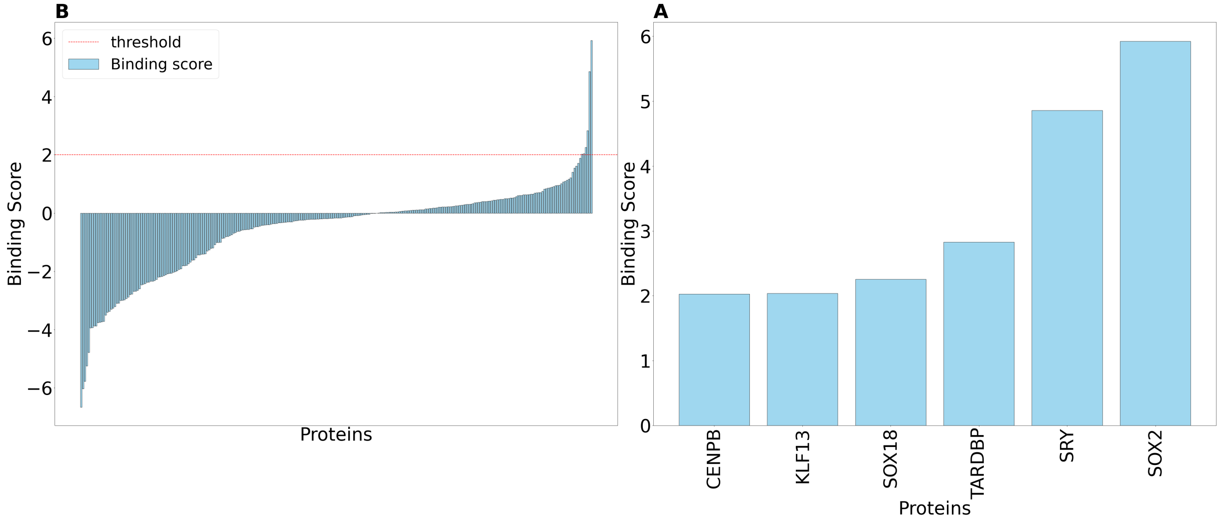


**Fig. 8. Specificity of Binding for Aptamer 2.**

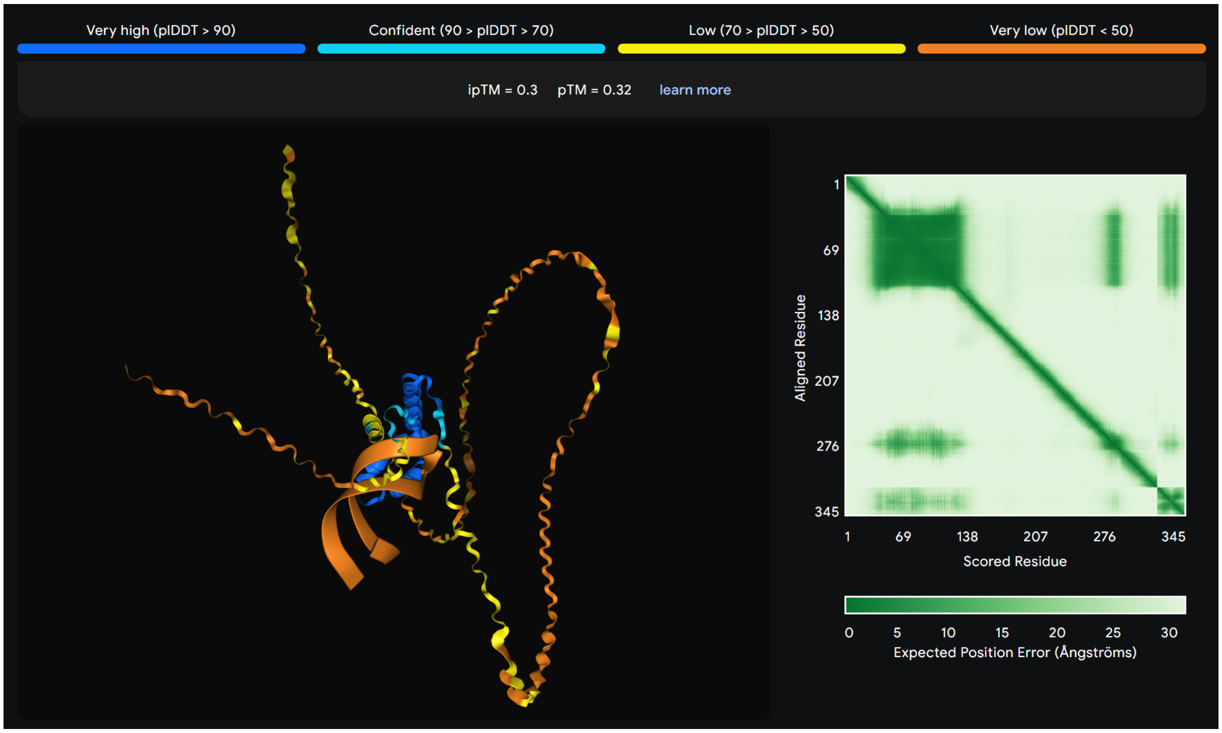


**Fig. 9.** Prediction of the structure of the full complex of Aptamer 1 and the SOX2 protein using AlphaFold 3.

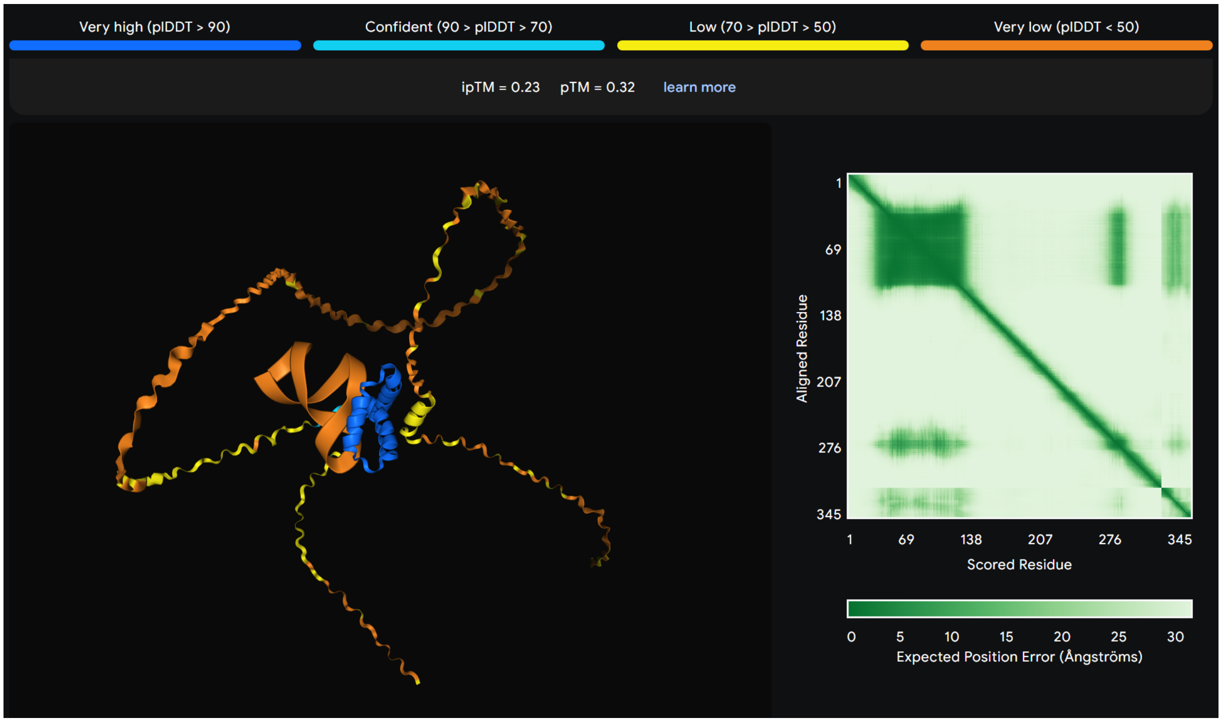


**Fig. 10.** Prediction of the structure of the full complex of Aptamer 2 and the SOX2 protein using AlphaFold 3.

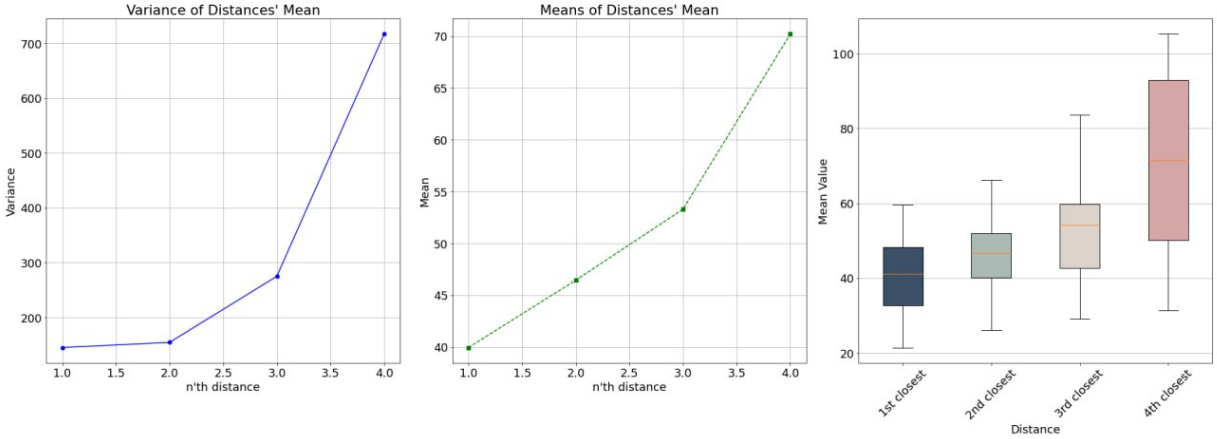


**Fig. 11. Analysis of the optimum number of proxy proteins.** The distances shown here are with respect to the target protein and in the ProtTrans space.

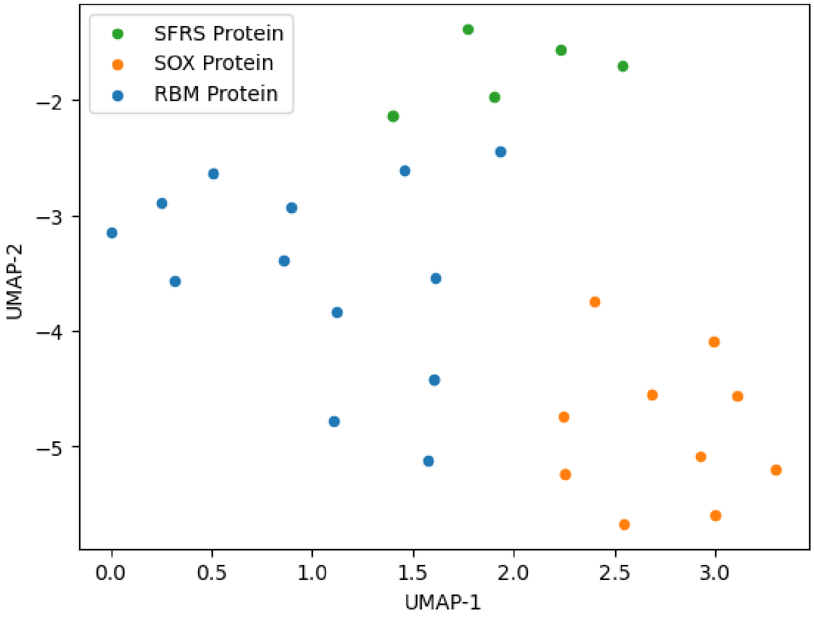


**Fig. 12. Reduced dimensions of the ProtTrans embeddings for the SOX2, SFRS, and RBM protein families using UMAP.**

### 2 Supplementary Notes

#### 2.1 Optimization settings for SOX4, SOX2 and RBM45

To perform the optimization, we selected SOX15, SOX14, and SOX7 proteins as the closest relatives for SOX2, and for SOX4, we selected SOX8, SOX7, and SOX10 proteins. For the RBM45 protein, we found RBM46, RBM41, and RBM4 proteins to be the closest relatives for which DeepBind models were available. We assumed that there are no DeepBind models for SOX4, SOX2, and RBM45 proteins, while DeepBind models are available for the selected relatives.

#### 2.2 Experimental Setup and Hyperparameter Optimization

The model is implemented using the Tensorflow 1 [1] library and in Python. We train and optimize the model on a Super-Micro Super-Server 4029GP-TRT with 2 Intel Xeon Gold 6140 Processors (2.3 GHz, 24.75 M cache), 256 GB RAM, 6 NVIDIA GeForce RTX 2080 Ti GPUs (11 GB, 352 bit), and 2 NVIDIA TITAN RTX GPUs (24 GB, 384 bit). We only use a single TITAN RTX GPU. The training takes around 86 hours for 200,000 epochs, and the optimization takes roughly 5 minutes for 3,000 iterations using GPU and 25 minutes using CPU.

#### 2.3 Optimization algorithm pseudocode

Here, we show the step-by-step process of optimization of the generated sequences.

**Algorithm 1:** Optimization of Z with the overall architecture

**Result:** RNA sequences with high binding preference to the targeted protein

$\theta \longleftarrow$ parameters of the trained GAN on any large RNA dataset. ;

$f_e(P) \longleftarrow$ protein embedding function, which outputs the embedding vector of its input amino acid sequence. ;

$\{F_i(R)\} \longleftarrow$ set of all available target feature assessment models. ;

$Z = Z_0 \longleftarrow$ initial latent noise sampled from standard normal.;

$P_t \longleftarrow$ amino acid sequence of the target protein.;

$N \longleftarrow$ number of closest proteins to be used.;

$x_t \longleftarrow f_e(P_t)$ ;

$\{x_i\} \longleftarrow f_e(\{P_i\})$ ;

$\{d_i\} \longleftarrow \{\|x_t - x_i\|^2\}$ ;

$\{d_i\}_{i=0}^{N} \longleftarrow \{d_i\}$ with $N$ of the minimum $d_i$ ;

$\{w_i\}_{i=0}^{N} \longleftarrow softmax(\{d_i\}_{i=0}^{N})$;

**for** $\hat{y_t}$ is not converged **do**

  $R \longleftarrow GAN_\theta(Z)$ ; // generated sequences

  $\{y_i\}_{i=0}^{N} \longleftarrow \{F_i(R)\}_{i=0}^{N}$ ; // assessment scores for $N$ closest proteins

  $\hat{y_t} \longleftarrow \sum_{i=0}^{N} y_i w_i$ ; // estimate of target protein binding score

  $\theta \longleftarrow \theta + _{\theta}^{\hat{y_t}}$ ; // update $\theta$ with SGA

**end**

$R^* \longleftarrow GAN_\theta(Z)$ ; // optimized sequences

#### 2.4 Hyperparameter optimization

The main hyperparameters of our model include the number of proxy proteins, the size of the latent dimension, the number of generator layers, and the number of discriminator layers. To find the best combination of parameters, we run a grid search on a reasonable range of parameters. The model is trained up to 200,000 generator iterations. Many of the instances of the model fail to learn and overfit, resulting in the validation loss going below the training loss, and the training loss begins to increase in some cases. In the cases that the mentioned scenario doesn't happen, we select the parameters where the $p$-values for the statistical tests are closer to desirable values. The latent dimension in RNAGEN is 25, and we present the number of layers in Supplementary Tables 3 and 4. As you can see in Supplementary Figure 9, we observe 3 to be the best number of proxy proteins for optimization. This is because as the distance to the target protein increases, the optimization performance will drop.

#### 3 Supplementary Tables

| **Name** | **Sequence** |
|---|---|
| T7 promoter | TTCTAATACGACTCACTATAGG |
| RNA aptamer 1 | UGGGAAGAAGAAUGAUAUUCUGUGUGUA |
| RNA aptamer 2 | GCUGGUGGUGAAUGACAUUGAUUUGAUCAA |
| SOX2 gene | ATGTACAACATGATGGAGACGGAGCTGAAGCCGCCGGGCCCGCAGCAAACTTCGGGGGGCG<br>GCGGCGGCAACTCCACCGCGGCGGCGGCCGGCGGCAACCAGAAAAACAGCCCGGACCGCG<br>TCAAGCGGCCCATGAATGCCTTCATGGTGTGGTCCCGCGGGCAGCGGCGCAAGATGGCCC<br>AGGAGAACCCCAAGATGCACAACTCGGAGATCAGCAAGCGCCTGGGCGCCGAGTGGAAAC<br>TTTTGTCGGAGACGGAGAAGCGGCCGTTCATCGACGAGGCTAAGCGGCTGCGAGCGCTGC<br>ACATGAAGGAGCACCCGGATTATAAATACCGGCCCCGGCGGAAAACCAAGACGCTCATGA<br>AGAAGGATAAGTACACGCTGCCCGGCGGGCTGCTGGCCCCCGGCGGCAATAGCATGGCGA<br>GCGGGGTCGGGGTGGGCGCCGGCCTGGGCGCGGGCGTGAACCAGCGCATGGACAGTTACG<br>CGCACATGAACGGCTGGAGCAACGGCAGCTACAGCATGATGCAGGACCAGCTGGGCTACC<br>CGCAGCACCCGGGCCTCAATGCGCACGGCGCAGCGCAGATGCAGCCCATGCACCGCTACG<br>ACGTGAGCGCCCTGCAGTACAACTCCATGACCAGCTCGCAGACCTACATGAACGGCTCGC<br>CCACCTACAGCATGTCCTACTCGCAGCAGGGCACCCCTGGCATGGCTCTTGGCTCCATGG<br>GTTCGGTGGTCAAGTCCGAGGCCAGCTCCAGCCCCCCTGTGGTTACCTCTTCCTCCCACT<br>CCAGGGCGCCCTGCCAGGCCGGGGACCTCCGGGACATGATCAGCATGTATCTCCCCGGCG<br>CCGAGGTGCCGGAACCCGCCGCCCCCAGCAGACTTCACATGTCCCAGCACTACCAGAGCG<br>GCCCGGTGCCCGGCACGGCCATTAACGGCACACTGCCCCTCTCACACATG |

**Table S1.** Sequences used in the in vitro experimentation.

| **Primer name** | **Sequence** |
|---|---|
| pET22B-PELB-Rev | GGCCATCGCCGGCTGG |
| pET22B-PELB-For | CTCGAGCACCACCACCACCA |
| primerDoA-142 | CTGCTCCTCGCTGCCCAGCCGGCGATGGCCATGTACAACATGATGGAGACGGAG |
| primerDoA-143 | ATCTCAGTGGTGGTGGTGGTGGTGCTCGAGCATGTGTGAGAGGGGCAGTG |

**Table S2.** Primers used in this study.

| Layer | Kernel size | Output shape |
|---|---|---|
| Input | - | 64 x 25 |
| Dense | - | 64 x 800 |
| Reshape | - | 64 x 32 x 25 |
| Convolution (1D) | 1 | 64 x 32 x 25 |
| Residual block (1D) | [5]x2 | 64 x 32 x 25 |
| Residual block (1D) | [5]x2 | 64 x 32 x 25 |
| Residual block (1D) | [5]x2 | 64 x 32 x 25 |
| Residual block (1D) | [5]x2 | 64 x 32 x 25 |
| Residual block (1D) | [5]x2 | 64 x 32 x 25 |
| Convolution (1D) | 1 | 64 x 32 x 5 |
| Softmax | - | 64 x 32 x 5 |

**Table S3.** The layers of the generator G in the WGAN.

| Layer | Kernel size | Output shape |
|---|---|---|
| Input | - | 64 x 32 x 5 |
| Convolution (1D) | 1 | 64 x 32 x 25 |
| Residual block (1D) | [5]x2 | 64 x 32 x 25 |
| Residual block (1D) | [5]x2 | 64 x 32 x 25 |
| Residual block (1D) | [5]x2 | 64 x 32 x 25 |
| Residual block (1D) | [5]x2 | 64 x 32 x 25 |
| Residual block (1D) | [5]x2 | 64 x 32 x 25 |
| Residual block (1D) | [5]x2 | 64 x 32 x 25 |
| Residual block (1D) | [5]x2 | 64 x 32 x 25 |
| Residual block (1D) | [5]x2 | 64 x 32 x 25 |
| Residual block (1D) | [5]x2 | 64 x 32 x 25 |
| Residual block (1D) | [5]x2 | 64 x 32 x 25 |
| Flatten | - | 64 x 800 |
| Dense | - | 64 x 1 |

**Table S4.** The layers of the discriminator D in the WGAN.
